# Ataxin-7 promotes Drosophila posterior follicle cell maturation by suppressing yorkie function

**DOI:** 10.1101/2022.08.03.502610

**Authors:** Hammed Badmos, Daimark Bennett

## Abstract

Ataxin-7 is a key component of the Spt-Ada-Gcn5-acetyltransferase (SAGA) chromatin-modifying complex that anchors Non-stop/USP22, a deubiquitinase, to the complex, thereby helping to coordinate SAGA’s different activities. Recently, we found that *non-stop* is required in the *Drosophila* ovary for expression of Hippo signalling pathway components *ex* and *mer*, and polarisation of the actin cytoskeleton during collective epithelial cell migration. Here we show that in addition to being required for collective migration, *Ataxin-7* plays an essential role in posterior follicle cells (PFCs) to control epithelial maturation and architecture, as well as body axis specification which depends on correct PFC differentiation. Loss of both the deubiquitinase and acetyltransferase modules of SAGA phenocopy loss of *Ataxin-7* in PFCs, demonstrating a redundant requirement for SAGA’s enzymatic modules. Strikingly, loss of *yorkie* completely suppressed effects of *Ataxin-7* loss-of-function in PFCs, indicating that the only essential function of *Ataxin-7* in PFCs is to suppress *yorkie* function. This may have broad relevance to the roles of SAGA and Ataxin-7 in development and disease.

## Introduction

The *Drosophila* ovary provides a tractable system for understanding the interplay between the genetic networks and cell interactions that control epithelial maturation and tissue development. The ovary consists of about 15-18 ovarioles, in which egg chambers at different stages of development, from the germarium through to the mature egg chamber, can be observed (King et al., 1956). Each egg chamber is comprised of a monolayer of somatically-derived follicle cells that surrounds 15 nurse cells and one oocyte that make up the germ-line. Communication to and from the follicular cells and the germline establishes patterning of both the somatic epithelium and the developing oocyte. Delta, a Notch ligand produced in the germline at around stage 6 of egg chamber development reprograms gene expression in follicle cells (Lopez-Schier and St Johnston, 2001; Ruohola et al., 1991) and signals the end of mitotic divisions in follicle cells as they begin to amplify their genome via endoreduplication (Deng et al., 2001; Lopez-Schier and St Johnston, 2001). Around this time, polar cells, located at the termini at each end of the egg chamber are specified in a Notch-dependent manner (Grammont and Irvine, 2001). Polar cells act as organising centres: at the anterior end, polar cells secrete the JAK/STAT ligand Unpaired to specify migratory border cells, stretched cells and centripetal cells during stages 7-9. At the posterior end, the TGFα-like ligand Grk, which is localised at the posterior pole of the oocyte by microtubules, signals to the overlying cells and they adopt the Posterior Follicle Cell fate (PFC) (Gonzalez-Reyes et al., 1995; Roth et al., 1995).

PFCs also receive signals from the Salvador-Warts-Hippo pathway, acting upstream of Notch, to suppress the transcriptional co-activator Yorkie, thereby controlling PFC maturation (Meignin et al., 2007; Polesello and Tapon, 2007; Yu et al., 2008). Once specified, the PFCs send a signal, which is yet to be identified, back to the oocyte, to repolarise the microtubule cytoskeleton. In turn, this directs the movement of the oocyte nucleus and polarising factors, such as *gurken* mRNA, to the dorsal-anterior corner of the oocyte, which is important for dorsal/ventral axis formation (Neuman-Silberberg and Schupbach, 1993; Neuman-Silberberg and Schupbach, 1996; Roth et al., 1995; Schupbach, 1987). Microtubule repolarisation also establishes the localisation of embryonic anterior/ posterior polarity determinants, (reviewed, Roth and Lynch, 2009). Consequently, maturation of the follicular epithelium is critical, not just for functional organisation of the soma, but also for establishing the body plan in the resulting embryo.

Developmentally-regulated patterns of gene expression, under epigenetic control, underpin the competence of cells to send and receive signals, which, in turn, determine the fate of cells within epithelia. Amongst many epigenetic regulators emerging as being important for developmentally-controlled gene expression is the ∼2-MDa Spt -Ada-Gcn5 acetyltransferase (SAGA) transcription coactivator complex, which consists of three essential modules: a histone acetyltransferase (HAT), a deubiquitinase (DUB) and a non-catalytic scaffold that provides structural stability to the complex (Chen and Dent, 2021; Soffers and Workman, 2020; Wang et al., 2020). Recent studies suggest that SAGA is responsible for the regulation of specific transcriptional states (Weake and Workman, 2012), which likely underpin roles of its various subunits in changes in cell polarity, cell specification and identity (Badmos et al., 2021; Koutelou et al., 2019). For instance, we recently found that Non-stop/USP22, the catalytic component of the DUB module, regulates F-actin and cell polarity of *Drosophila* border cells, in part by promoting the expression of Hippo signalling pathway components, *mer* and *ex* (Badmos et al., 2021). Hippo components function at border cell-border cell junctions to phosphorylate and inhibit the actin regulator Enabled (Lucas et al., 2013). Here we show that another SAGA component, Ataxin-7, also regulates F-actin polarity and border cell migration. Furthermore, these studies reveal a novel role for SAGA components in Notch-mediated maturation of the follicular epithelium and body axis formation. Notably, effects of loss of *Ataxin-7* function in PFCs can be completely suppressed by loss of *yorkie*, indicating that *Ataxin-7* functions to maintain a Hippo signalling-dependent cell differentiation programme in PFCs.

## Results

### *Ataxin-7* is required for border cell migration

We recently identified *non-stop* in a screen for DUBs involved in border cell migration (Badmos et al., 2021) but the involvement of other DUB module components, such as *Ataxin-7* were not known. To address this issue, we used a null *Ataxin-7* allele (*Atxn7*^*HO3*^), which has a 1949bp deletion from exon 1 to exon 2 including the translation start codon and fails to produce detectable protein (Li et al., 2017). Using MARCM (Lee and Luo, 2001), we generated *Ataxin-7* loss-of-function clones of border cells labelled with GFP or RFP (**Fig.1**). We examined the effect on migration at stage 10 of egg chamber development, when border cells normally have completed their migration to the oocyte. All clusters analysed had greater than 50% mutant cells per cluster. Notably, less than 1% of *Atxn7*^*HO3*^ clusters completed their migration, and there was a significant reduction in mean migration compared to wild type controls (mean ±SEM was 19.0 ±4.9% n=33 clusters/replicate for *Atxn7*^*HO3*^, compared to 96.6 ±0.8% n=30 clusters/replicate for controls) (**Fig.1A-C**). For border cell clusters to migrate they need to acquire highly polarised actin protrusions at their leading edge at the start of migration (Bianco et al., 2007). Factin then becomes predominantly located around the cortex of wild type clusters. In contrast, we found that there was a shift in the distribution of F-actin toward the interior border cell-border cell junctions in *Ataxin-7*^*HO3*^ clones (**Fig.1A-B,D**). Similar to *non-stop* loss-of-function, we also observed a 2.2 fold increase in the number of border cells recruited into *Atxn7*^*HO3*^ clusters (mean number for *Atxn7*^*HO3*^ clusters was 12.8 border cells, n=33 clusters/replicate compared to 6.8 for wild type controls, n=30 clusters/replicate, *P*=0.004)(**Fig.1E-F,G**). Taken together, *Ataxin-7* phenocopies *non-stop* loss of function in migrating border cells.

**Fig.1.**
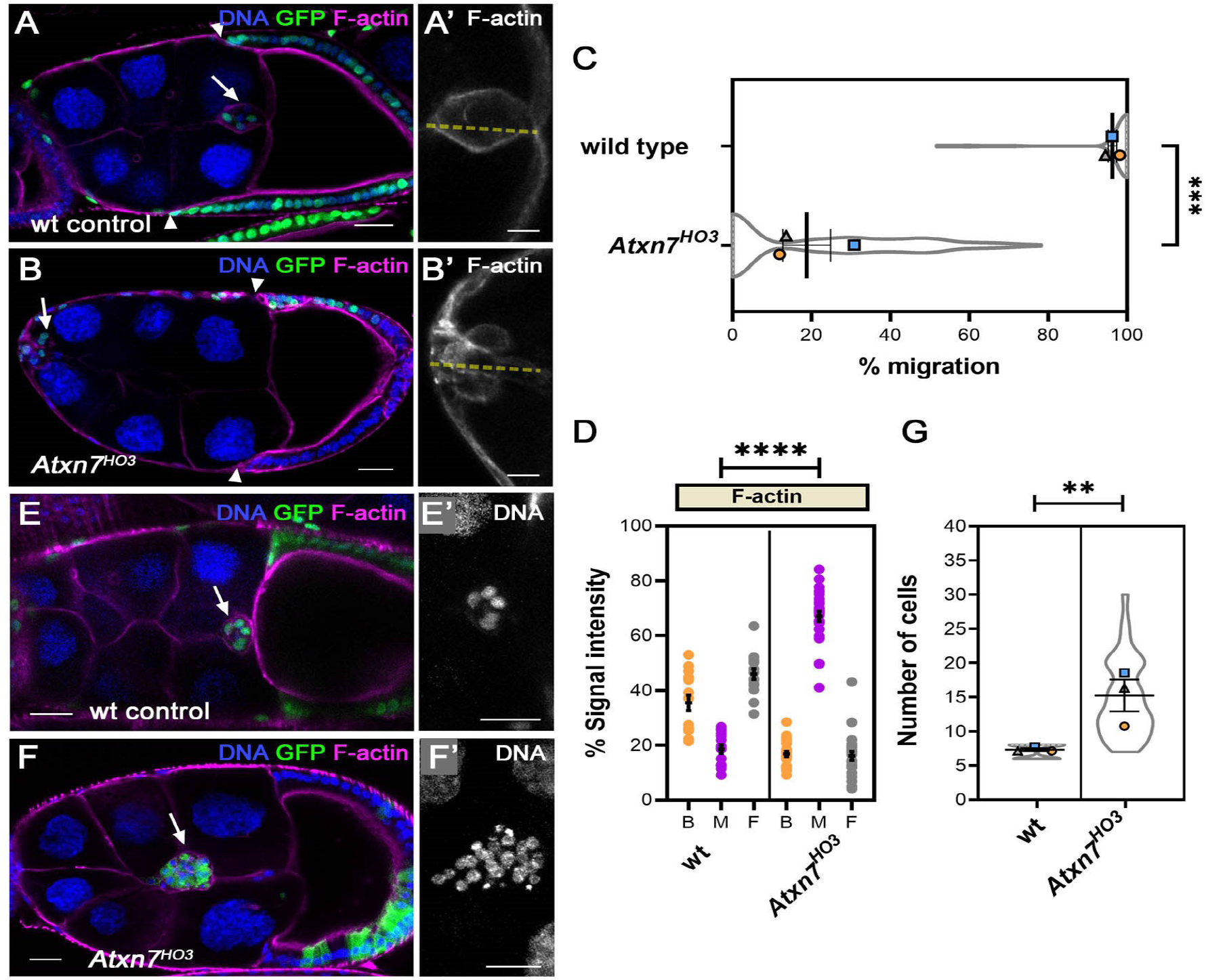
Ataxin-7 is required for border cell migration and number. **A-B**, Confocal images of late stage 9 egg chambers with GFP-labelled wildtype (wt) or *Atxn7*^*HO3*^ cells (green) generated with MARCM. F-actin is visualised with phalloidin staining (magenta) and nuclei are labelled with TOPRO-3 (blue). In control egg chambers (**A**), border cell (arrows) migrate normally in concert with centripetal follicle cells (arrowheads), whereas *Atxn7*^*HO3*^ border cell cluster shows strongly abrogated migration (**B**). Inset **A’** shows normal polarisation of actin cytoskeleton around the cortex of the wild type migrating cluster (grayscale). Inset **B’** shows accumulation of F-actin into junctions between *Atxn7*^*HO3*^ outer border cells (grayscale). Dotted line shows position of line scans taken for quantitation of Factin polarity. **C**, Plots showing mean % border cell migration from the anterior of the egg chamber to the nurse cell/oocyte boundary, ± SEM derived from means of three replicates, superimposed on violin plots of migration measurements for each indicated genotype. ***, P < 0.001, one -way ANOVA. **D**, Dot plots of area under curve derived from line scans through the cluster, as (Badmos et al., 2021), showing F-actin signal intensity at the back, middle, and front (B,M,F) of the BC cluster. Mean ±SEM is derived from three replicates, n>5 egg chambers/replicate, showing a consistent defect in F-actin polarization in *Atxn7*^*HO3*^ clusters. ****, P < 0.0001, one-way ANOVA. **E-F**, Confocal images of stage 10 egg chambers with GFP-labelled clones (green), phalloidin-stained F-actin (magenta) and TOPRO-3-labelled nuclei (blue). Compared to wild type controls, which typically contain up to 8 cells/cluster (**E, E’**), *Atxn7*^*HO3*^ border cell clusters (**F, F’**) show as many as 30 cells. (**G**) Violin plot with means ± SEM (n=3) showing a significant increase in border cell numbers in *Atxn7*^*HO3*^ clusters compared to controls. Bars, 25μm.

### *Ataxin-7* is required in follicle cells to control cell proliferation and polarity

In wild type egg chambers a single layer of follicle cells surrounds the germ cells, with PFCs adopting a monolayered columnar epithelium at stage 10 (**Fig.2A,A’**). In contrast, *Ataxin-7*^*HO3*^ cells at the posterior end of the egg formed a bi- or multi-layered epithelium (61 ±6.9% of egg chambers, n=57 egg chamber/replicate) (**Fig.2B-C**). We also observed this at a lower frequency at the anterior end (51 ±2.9% of egg chambers), but almost never in the lateral regions of the egg chamber (1.3 ±0.67% of egg chambers) (**Fig. 2D**). Notably, at a lower frequency, we also observed non-uniform multilayering of *Ataxin-7*^*HO3*^ follicle cells (**Fig. 2C,D**). Time-lapse imaging revealed that non-uniform bulging of the epithelium was accompanied by localised invasion of some cells into the centre of the egg chamber (**Fig. 2C, video 1** and **2**). This mostly occurred at the posterior (38 ±7.5%) and anterior (33 ±6.0%) ends of the egg chamber and only very rarely in the lateral region of the epithelium (1.3 ±1.3%) (**Fig. 2D**).

**Fig.2.**
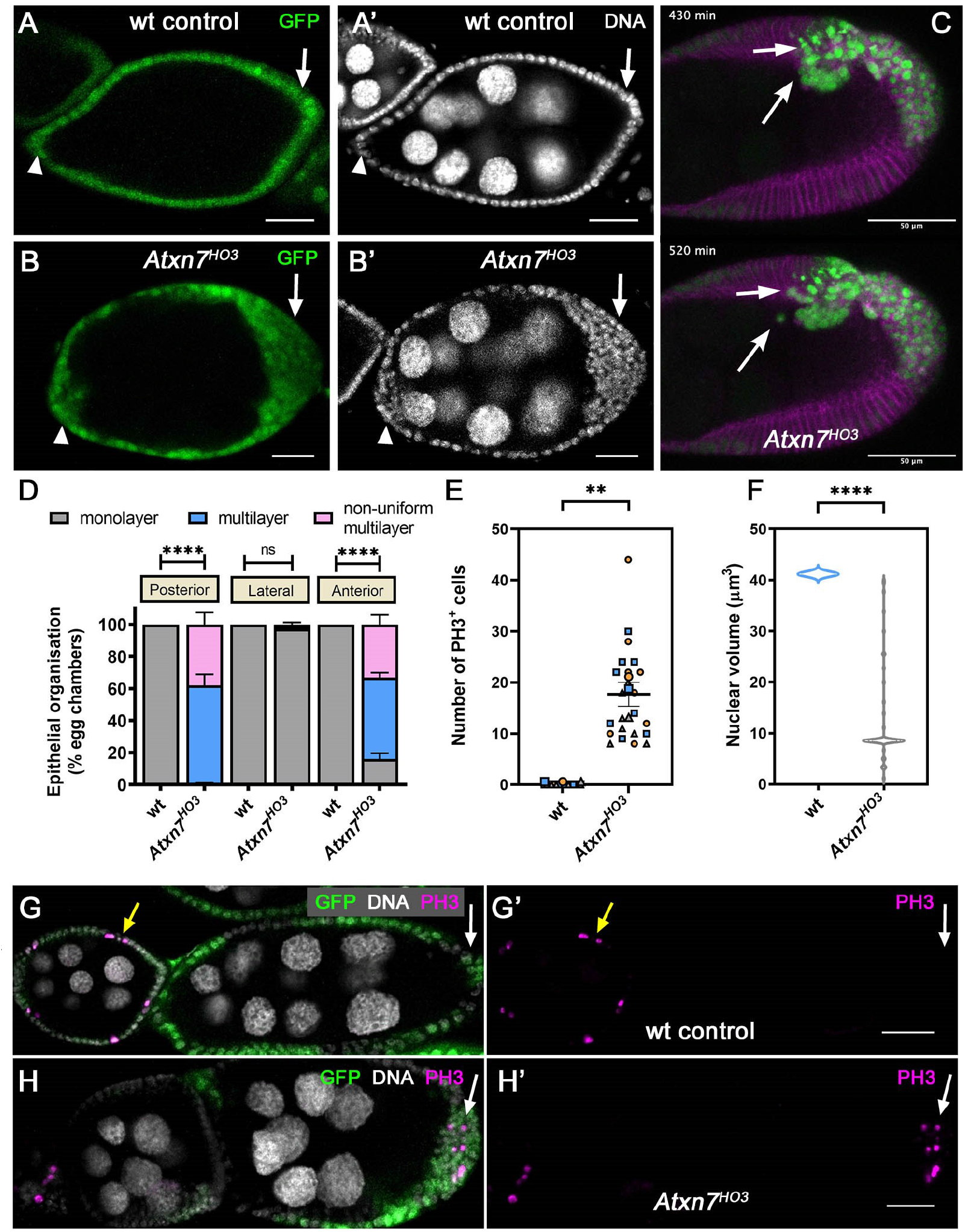
*Ataxin-7* is required for normal follicle cell epithelial architecture. **A-B**, Confocal sections of stage 7 egg chambers with GFP-labelled clonal cells (green) generated with MARCM. All nuclei were labelled with TOPRO-3 (greyscale). Follicular epithelium (arrows) shows normal monolayered architecture (**A, A’**) whereas *Atxn7*^*HO3*^ shows an abnormal architecture that is multi-layered (**B, B’**). **C**, Still images from timelapse (video 2), showing non-uniform multilayering of GFP-labelled *Atxn7*^*HO3*^ cells and movement of a subset of these towards and into the interior of the egg chamber (see arrows). Membrane is labelled with FM4-64 dye (magenta). (**D**) Stacked bar chart showing quantification of epithelial architecture in wt and *Atxn7*^*HO3*^ clones located at the anterior, lateral or posterior regions of the egg chamber. (**E**) Superplot showing quantitation of the number of PH3 positive cells after stage 6 of mitosis phase during egg development. Means ± SEM (n=3). **, P < 0.01, one-way ANOVA. (**F**) Violin plot showing nuclear volume of follicle cells visualised with TOPRO-3 staining. ****, P < 0.0001, one-way ANOVA. **G-H**, Confocal sections of stage 6-8 egg chambers with GFP-labelled clonal cells (green), TOPRO-3-labelled nuclei (greyscale) and mitotic nuclei labelled with anti-phosphohistone H3, PH3 (magenta). In controls, PH3 is only ever seen in epithelia of early stage egg chambers (yellow arrow), whereas PH3 positive nuclei are frequently observed in multi-layered *Atxn7*^*HO3*^ epithelia at later stages (compare PFCs in control and *Atxn7*^*HO3*^ egg chambers indicated with white arrows). Bars, 25μm.

To determine whether multilayering phenotypes were due to overproliferation, which can result from a failure to switch from a mitotic cell cycle to an endocycle at stage 6 of oogenesis, we labelled cells with the mitotic marker Phospho-Histone H3 (PH3). After stage 6, we observed an absence of PH3 staining in wild type egg chambers, but in *Atxn7*^*HO3*^ clones we routinely saw PH3 staining at the anterior and posterior ends of egg chambers up to stage 10 (**Fig.2E-H**). Analysis of the size of nuclei in the follicular epithelium also revealed a 3.4-fold reduction in mean nuclear volume (mean volume was 41.2±0 μm^3^ for controls and 12.2±1.35 μm^3^ for *Atxn7*^*HO3*^ clones), consistent with a failure to undergo endoreplication (**Fig.2F**). Taken together, these data suggest that *Ataxin-7* controls proliferation of follicle cells located at the ends of the egg chamber.

Multilayering and invasion of follicle cells into the oocyte has been described for *discs-large* (*dlg*) loss-of-function mutations (Goode and Perrimon, 1997), prompting us to examine effects of *Atxn-7* loss-of-function on cell polarity. Notably, we observed loss of apical Crumbs (Crb) and atypical protein kinase C (aPKC) (**Fig.3A-E**). Crb and aPKC staining at the apical domain was reduced 4.9-fold (n≥15, *P<*0.0001, unpaired t-test) and 9.2-fold (n≥8, *P<*0.0001, unpaired t-test), respectively (**Fig.3C**). Polarisation of basolateral Dlg and apical/lateral Armadillo/β-catenin (Arm) was also disrupted, with both being spread around the cortex of cells, which appeared some-what rounded having lost their columnar appearance (**Fig.3F-I**). These results show that *Ataxin-7* is required for the normal localisation of cell polarity determinants and normal epithelial architecture.

**Fig.3.**
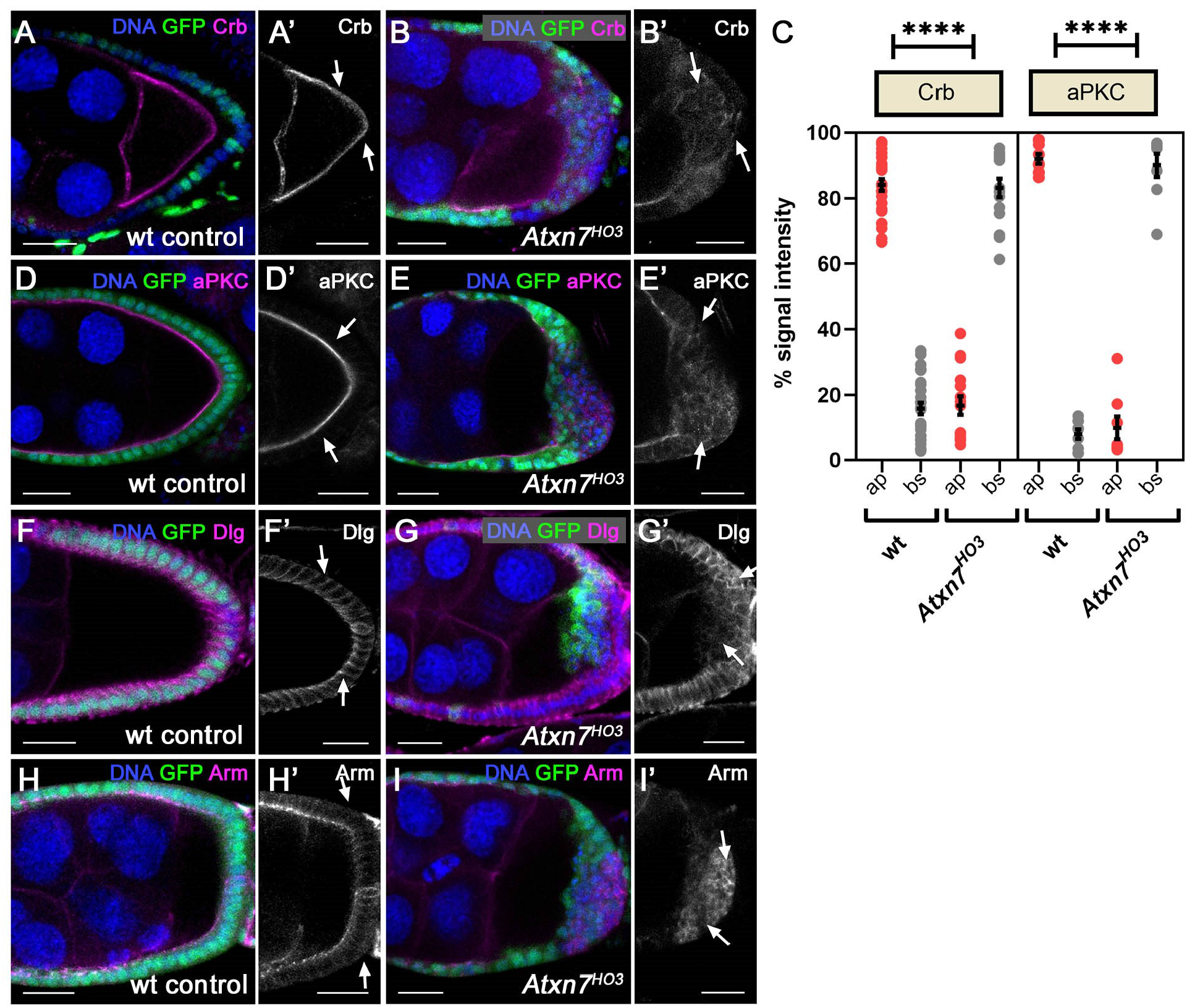
*Ataxin-7* loss of function disrupts membrane polarity. **A-B**, Confocal images of egg chambers with GFP-labelled clones (green) stained with antibodies against Crb (magenta) and TOPRO-3 to label nuclei (blue). In wild type egg chambers, Crb is localised at the apical face of posterior follicle cells (**A, A’** see arrows). This distribution is lost in *Atxn7*^*HO3*^ cells (**B, B’** see arrows). **C**, Dot plots of area under curve derived from line scans of signal intensity for Crb and aPKC across apical (ap) and basal (bs) sides of the posterior follicular epithelium taken from egg chambers with wildtype or *Atxn7*^*HO3*^ clones. Mean ± SEM of three replicates. ****, P < 0.0001, one-way ANOVA. Apical Crb and aPKC staining is lost in *Atxn7*^*HO3*^ clones. **D-I**, Confocal images of egg chambers with GFP-labelled clones (green) stained with antibodies against aPKC (**D-E**), Dlg (**F-G**), Arm (**H-I**) (magenta) and TOPRO-3 to label nuclei (blue). Basolateral Dlg (F) and apical/lateral Arm (H) was also disrupted in *Atxn7*^*HO3*^ clones, with both being spread around the cortex of *Atxn7*^*HO3*^ cells, which lose their columnar appearance (see arrows). Bars, 25μm.

### *Ataxin-7* is required for normal Notch-dependent PFC maturation

Failure of *Ataxin-7* mutant PFCs cells to exit mitosis suggests that they are stuck in an immature state and their differentiation is impaired. To explore this, we examined effects on Notch signalling, which facilitates a transcriptional programme at mid-oogenesis triggering the mitotic-endocycle switch and subsequent PFC maturation (Deng et al., 2001; Lopez-Schier and St Johnston, 2001). In *Ataxin-7* mutant PFCs we observed a disruption of Notch localisation, possibly as a consequence of impaired cell polarity (Li et al., 2009) (**Fig.4A-B**). To assess effects on downstream targets of Notch, we analysed the expression of Cut, a homeobox protein, and Cyclin B, a mitotic cyclin, which are down-regulated by Notch signalling from stage 7 onwards (Sun and Deng, 2005). Consistent with disrupted Notch signalling we found that Cut and Cyclin B showed continued expression in *Ataxin-7*^*HO3*^ clones at later stages of egg chamber development (**Fig.4C-F**). In wild type controls, expression of Cut and Cyclin B ceased at stage 6 (n=16 and n=10, egg chambers respectively), but both Cut and Cyclin B staining persisted in *Ataxin-7*^*HO3*^ PFC clones until stage 10 of egg chamber development (n=21, n=9 respectively).

**Fig.4.**
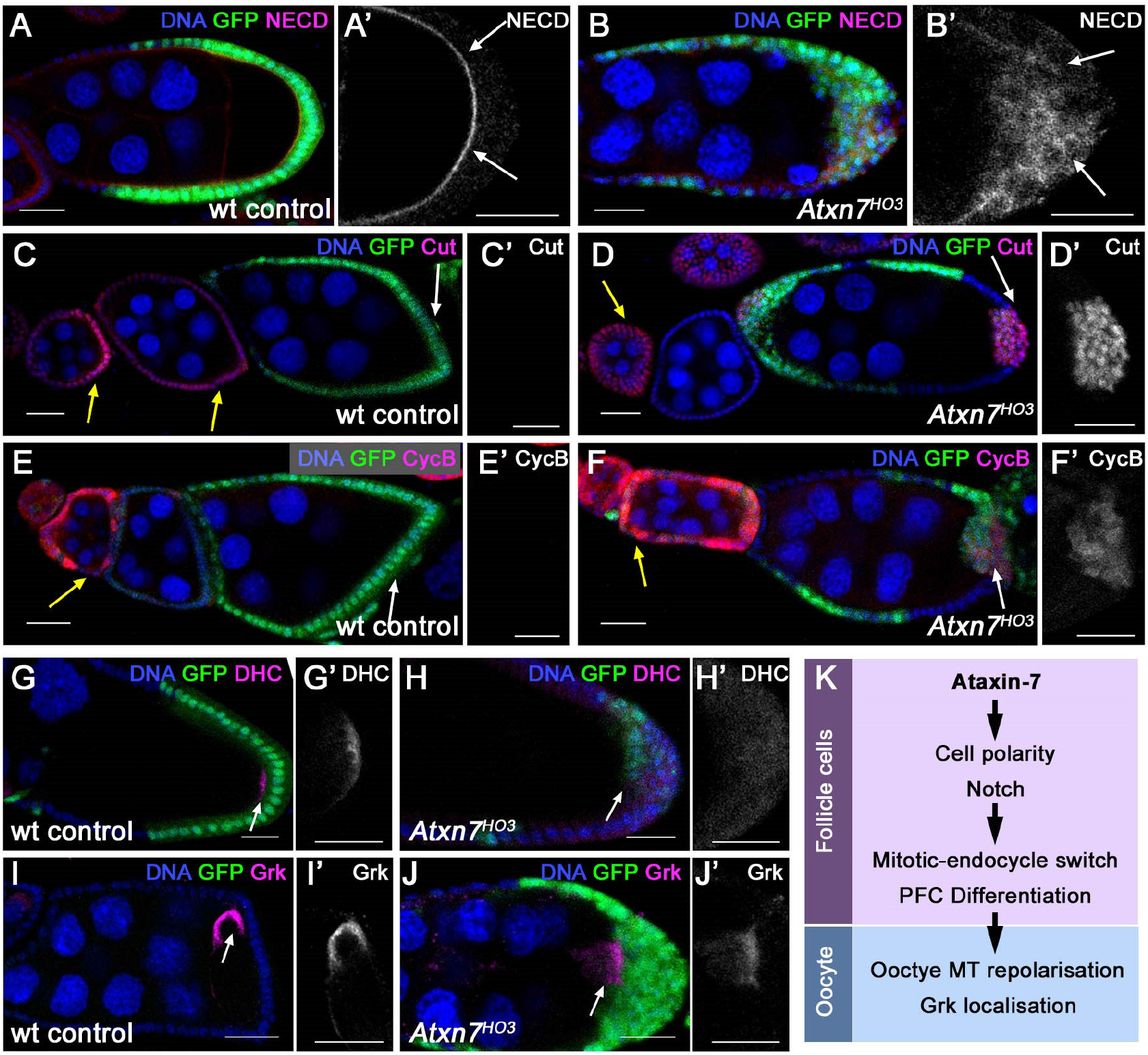
*Ataxin-7* loss of function disrupts PFC differentiation and oocyte polarity. **A-J**, Confocal images of egg chambers with GFP-labelled wild type or *Atxn7*^*HO3*^ cells (green) and TOPRO-3 labelled nuclei (blue) stained with antibodies (magenta, and grayscale in magnified images) against: (**A-B**) Notch extracellular domain (NECD); (**C-D**) Cut; (**E-F**) Cyclin B (CycB); (**G-H)** Dynamin Heavy Chain, (DHC); and (**I-J**) Gurken (Grk). Bars, 25μm. Wild type follicular epithelium shows normal apical NECD staining (**A-A’**, arrows), whereas in *Atxn7*^*HO3*^ cells, NECD is mis-localised (**B-B’**). **C-D**, Anti-Cut staining shows that wild type follicle cells show presence of Cut before stage 6 of egg development (yellow arrow) and absence of Cut after stage 6 of egg development (white arrow) (**C**). In contrast, *Atxn7*^*HO3*^ cells show presence of strong Cut expression after stage 6 of egg development (white arrow) (**D**). Similarly, wildtype follicular epithelium shows presence of CycB before stage 6 of egg development (yellow arrow) and absence of CycB after stage 6 (white arrow) (**E**), whereas *Atxn7*^*HO3*^ cells shows abnormal presence of CycB after stage 6 (white arrow) (**F**). **G-H**, Anti-Dynein heavy chain staining (red and grayscale) shows wildtype egg chambers have a normal Dhc localisation at the cortex of the posterior end of the oocyte (**G,G’**) whereas egg chambers with *Atxn7*^*HO3*^ follicle cell clones show a weak diffuse distribution of Dhc in the oocyte (**H, H’**). **I-J**, Correspondingly, in wildtype egg chambers there is a normal distribution of Grk to the anterior dorsal corner of the oocyte (arrow) (**I,I’**) whereas egg chambers with *Atxn7*^*HO3*^ PFCs, show an abnormal Grk localisation at the posterior (arrow) (**J,J’**). **K**, Schematic illustration summarising events dependent on *Atxn7* function in follicle cells.

### *Ataxin-7* is required for oocyte polarity

Correct specification and maturation of the PFCs is necessary for the ability of PFCs to signal to the oocyte and trigger the reorganisation of the microtubule cytoskeleton, which subsequently leads to the polarisation of the AP and DV axes. This prompted us to examine whether egg chambers lacking *Ataxin-7* in PFCs resulted in a failure to repolarize the oocyte microtubule cytoskeleton and polarity determinants. To do this, we visualised the localisation of Dynein heavy chain (Dhc), a microtubule motor that is normally found at posterior pole of the oocyte at stage 9. We found that Dhc was no longer tightly localised at the posterior of the oocyte in stage 9 egg chambers harbouring *Ataxin-7*^*HO3*^ posterior follicle cell clones, and was instead diffusely distributed (n=7) (**Fig.4G-H**). Dynein heavy chain (Dhc) is required for relocalisation of the oocyte nucleus, along with *gurken* mRNA and Gurken protein, to the anterior-dorsal region of the oocyte, which establishes DV polarity of the oocyte and the embryo (Januschke et al., 2002; Roth, 2003; St Johnston, 2005). Unlike in wild type controls (n=8), in stage 8-10 egg chambers with *Ataxin-7*^*HO3*^ posterior follicle cell clones, Grk was still anchored at the posterior of the oocyte, along with the oocyte nucleus (n=7) (**Fig.4I-J**). Taken together, we conclude that egg chambers with PFCs lacking *Ataxin-7* fail to signal to the oocyte, most probably due to a defect in their maturation, to induce correct repolarisation and axis specification of the oocyte (**Fig.4K**).

### Loss of both *Ada2b* and *Not*, but not either alone, phenocopies loss of *Ataxin-7*

The SAGA complex consists of two enzymatic modules: a histone acetyltransferase (HAT) and deubiquitinase (DUB) (Weake and Workman, 2012). To determine the contribution of these modules to the maturation of the follicular epithelium, we analysed the effect of *non-stop* (the enzymatic component of the DUB) and *ada2b* (an essential HAT component) loss-of-function. Knock-down of *non-stop* (**Fig.5A-B**) or mutation of *ada2b* in PFCs (**Fig.5C-D**) did not result in any detectable epithelial defect and was not accompanied by ectopic expression of Cut or mislocalisation of Crb (n=25 and n=14, respectively). However, loss-of-function of both *non-stop* and *ada2b* together (*non-stop* knockdown in *ada2b*^*1*^ clones) resulted in bilayering of the posterior follicular epithelium, ectopic expression of Cut and mislocalisation of Crb, phenocopying the effect of *Ataxin-7*^*HO3*^ (n=7), albeit more weakly (**Fig.5E-F**).

**Fig.5.**
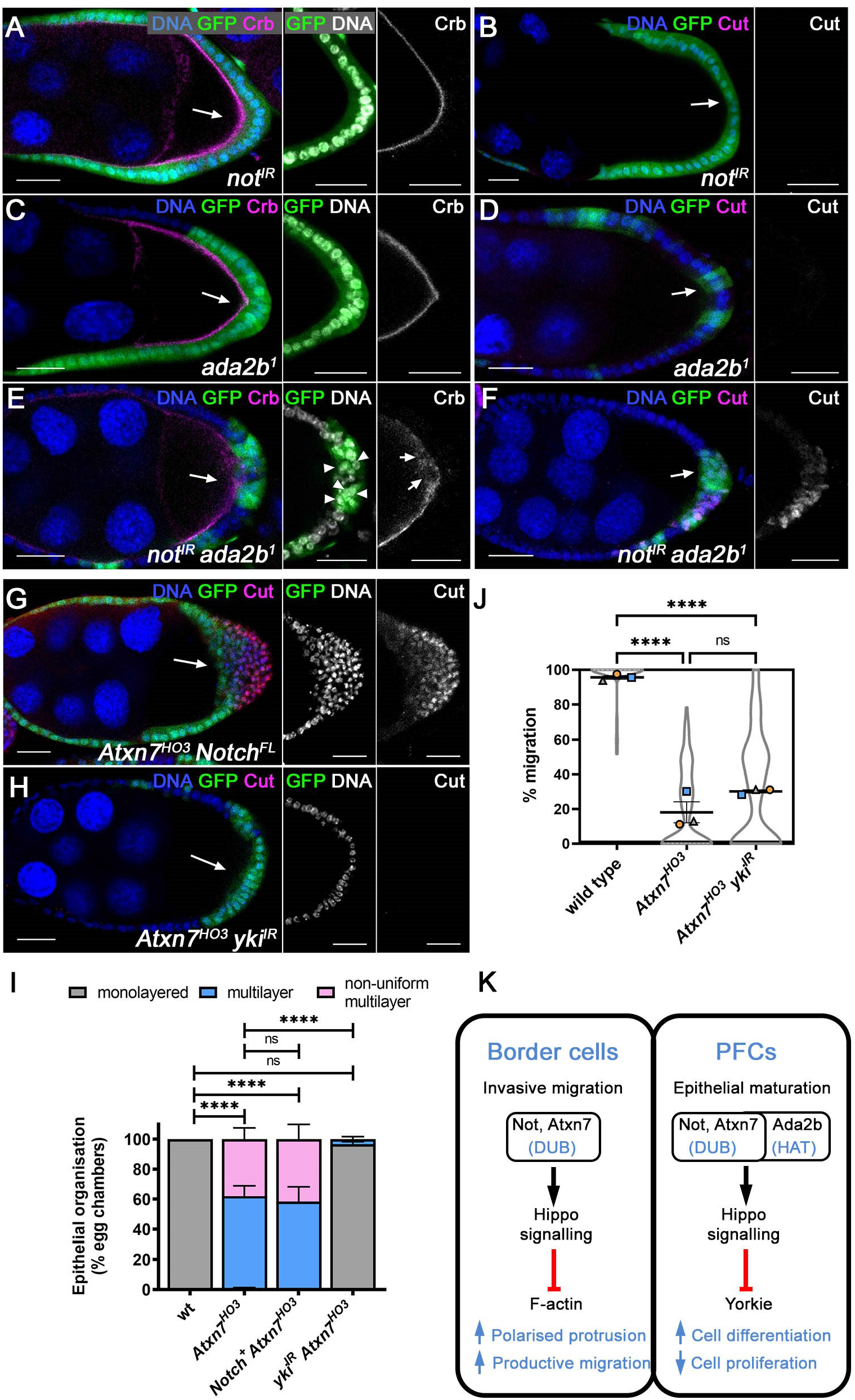
Ataxin-7 is phenocopied by ada2b/not and rescued by yki loss of function. **A-H**, Confocal images of egg chambers with GFP-labelled clones of the indicated genotypes (green) stained with TOPRO-3 to label nuclei (blue) and with antibodies against Crb (**A,C,E** in magenta and grayscale) or Cut (**B,D,F-H** in magenta and grayscale). Arrows indicate PFCs. Bars, 25μm. The follicular epithelium shows normal architecture, apical Crb and absence of Cut staining in *non-stop* knockdown (*not*^*IR*^, **A,B**) or *ada2b*^*1*^ clones (**C,D**). However, multilayering of the epithelium (arrowheads) and ectopic Cut are clearly evident when *not*^*IR*^ and *ada2b*^*1*^ are combined (**E,F**). Overexpression of full-length Notch (*Notch*^*FL*^) failed to rescue *Atxn7*^*HO3*^-induced epithelial multilayering or ectopic Cut expression (**G**), whereas *yki* knockdown (*yki*^*IR*^) restored a monolayered epithelium lacking Cut expression in *Atxn7*^*HO3*^ clones (**H**). **I**, Stacked bar chart showing quantification of epithelial architecture of posterior follicle cell clones. **** P<0.0001; ns, not significant, one-way ANOVA. **J**, Plots showing mean % border cell migration from the anterior of the egg chamber to the nurse cell/oocyte boundary, ± SEM derived from means of three replicates, superimposed on violin plots of migration measurements for each indicated genotype. *** P<0.001; ns, not significant, one -way ANOVA. *yki*^*IR*^ does not significantly rescue border cell migration of *Atxn7*^*HO3*^ clones. **K**, Illustration summarising context-dependent ability of DUB and HAT components to support the function of Hippo signalling outputs in border cells and PFCs, as reported here and in (Badmos et al., 2021). In border cells, the Hippo complex suppresses Enableddriven F-actin formation at border cellborder cell junctions, which supports polarised protrusions and productive migration. In PFCs, Hippo signalling suppresses the transcriptional regulator Yorkie, thereby promoting differentiation and preventing cell proliferation. Various Hippo signalling components are transcriptional targets of Not and Ada2b putting SAGA upstream of Hippo signalling. We have not ruled out the possibility that SAGA also has Hippo-independent effects on Yorkie function in PFCs.

### The essential role of *Ataxin-7* in PFCs is to promote hippo signalling

In PFCs, the Hippo pathway functions to prevent the transcriptional co-activator Yorkie from driving a pro-proliferative transcriptional programme (Meignin et al., 2007; Polesello and Tapon, 2007; Yu et al., 2008). Numerous Hippo signalling pathway components are potential transcriptional targets of both Ada2b and Not in *Drosophila* (Badmos et al., 2021). This prompted us to test whether defects in epithelial maturation we observed in *Ataxin-7* mutant clones could be alleviated by modulating Notch or Hippo signalling. Overexpression of Notch failed to rescue *Ataxin-7* follicular epithelial phenotypes, whereas *yorkie* knockdown strongly rescued Cut expression and epithelial multilayering induced in *Ataxin-7* mutant follicle cells (**Fig.5H-I**). Notably, *yorkie* knockdown did not rescue *Ataxin-7*^*HO3*^ border cell phenotypes, including defective migration (**Fig.5J**), consistent with previous findings indicating that Hippo signalling functions in a *yorkie*-independent manner in border cells (Badmos et al., 2021; Lin et al., 2014; Lucas et al., 2013).

## Discussion

### Ataxin-7 and the enzymatic modules of SAGA are required for follicular epithelial maturation

Here we find that loss of *Ataxin-7* in cell clones disrupted the maturation of the follicular epithelium, leading to multilayering and an invasive phenotype at the anterior and posterior ends of the egg chamber. Defective maturation of PFCs was characterised by loss of epithelial polarity, disruption of Notch signalling and impaired cell differentiation. This was accompanied by follicle cell overproliferation, resulting in loss of PFC-mediated signalling and oocyte mispolarisation. These effects were partially phenocopied when we targeted both *not* and *ada2b*, suggesting HAT and DUB modules play a redundant role in follicle cell development. This redundancy is likely to be exhibited at the level of target genes, where the combined activities of histone acetylation and deubiquitination contribute to destabilisation of nucleosomes, thereby facilitating the passage of RNA polymerase II along genes that are targeted by SAGA (Batta et al., 2011). Interestingly, the non

-redundant role of Ataxin-7 that we report here in PFCs suggests that it may coordinate SAGA’s HAT and DUB activities, consistent with previous reports that HAT activity is dependent on the composition of HAT-associated protein complexes (Balasubramanian et al., 2002; Grant et al., 1999), and that loss of Ataxin-7 effects both H2B ubiquitination and H3K9 acetylation levels in flies (Mohan et al., 2014).

### SAGA components promote hippo signalling in different contexts

Together with our previous study of not/USP22 (Badmos et al., 2021), our findings here reveal different requirements for SAGA components in cells derived from the same epithelium. *Ataxin-7*, like *not*, is required for border cell migration, whereas *ada2b*, is dispensable for border cell migration but contributes to the maturation of PFCs. A different requirement for HAT/DUB components in different cell types might reflect functional differences in acetylation/ deubiquitination at specific promoters or gene bodies, or point to SAGA-independent roles for the DUB module, possibly via interactions with cell type-specific co-activators independent of SAGA. However, in both contexts that we have examined in the ovary, a critical role is to promote Hippo signalling outputs: in border cells, Not/ USP22 is necessary for expression of *mer* and *ex* to ensure F-actin polarity; in PFCs, Ataxin-7 restrains *yorkie* function to maintain a Hippo signalling-dependent cell differentiation programme (**Fig.5K**). Both Ada2b and Not are found associated with promoters of multiple Hippo pathway components in *Drosophila* embryos, raising the possibility that several Hippo pathway components may rely on DUB/HAT for their expression in PFCs. Current efforts are directed at exploring single-cell profiling technologies that may allow us to address this issue. Notably, *yki* knockdown completely suppressed the effects of *Ataxin-7* loss-of-function, including invasion of follicle cells into the central regions of the egg chamber. Invasion of follicle cells is not seen upon loss of *Warts* (Zhao et al., 2008), which directly regulates Yki activity. This suggests that invasion is contingent on a failure to suppress Yki, but requires other Hippo-independent events driven by loss of *Ataxin-7*. Such events might include a disruption of cell polarity determinants such as Dlg, which has been reported to give rise to similar follicle cell invasion phenotypes to the ones we have observed (Goode and Perrimon, 1997).

### Relevance to development and disease

Transitions in cell behaviour, from proliferation to differentiation and the acquisition of cell motility, underpin important phases of tissue and organ development, and are milestones in the progression of many diseases. Our findings identify SAGA components as important regulators of these transitions in the *Drosophila* ovary. Several studies point to a wider role for Ataxin-7 and SAGA in different developmental contexts and in disease states such as tumourigenesis (Montero-Conde et al., 2017). An investigation into USP22/SAGA-dependent regulation of Hippo signalling in these contexts may be warranted in the light of our findings.

## Methods

### Drosophila stocks and genetics

Flies were raised and crossed at 25°C according to standard procedures.

The following fly lines were obtained from the Bloomington Drosophila Stock Center: *FRT40A (*BL1816), *slbo-Gal4, UAS-GFP* (BL6458, Montell Lab), *UAS-Notch*^*FL*^ (BL26820), or the Vienna Drosophila Resource Centre: *UAS-not*^*IR*^ (GD11236), *UAS-yki*^*IR*^ (GD11187). *Ataxin-7*^*HO3*^ *FRT40A* and *FRT82B ada2b*^*1*^ were gifts from Jerry Workman (Li et al., 2017).

We used the following strains to make positively-marked GFP-labelled clones by Mosaic Analysis with a Repressible Cell Marker (MARCM) (Lee and Luo, 2001).

*hsFLP; tub-Gal80, FRT40A; Act>CD2>Gal4,UAS-GFP/ SM5-TM6B* (generated from BL5192, *Act>CD2>Gal4, UAS-GFP and hsFLP; If/SM5-TM6B)*

*hsFLP; Act>y>Gal4,UAS-GFP;FRT82B tubGAL80/SM5-TM6B* (generated from BL44408, *Act>y>Gal4, UAS-GFP and hsFLP; If/SM5-TM6B*).

*hsFLP; Act>y>Gal4,UAS-GFP* (generated from *Act>y>Gal4, UAS-GFP and hsFLP; If/SM5-TM6B*).

To obtain mitotic clones in follicle cells, progeny of the right genotypes were heat shocked twice a day for 1 hour each with at least 5 hr intervals between treatments, from pupae to adult at 37°C. Newly enclosed adults (2-3 d old) were fattened for 2 d on yeast paste. Information on these strains is also available at http://www.flybase.org.

### Immunofluorescent imaging

Ovaries were dissected in PBS (Phosphate buffer saline) and fixed with 3.7% paraformaldehyde in PBS. The ovaries were washed with PBST (1X PBS, 0.2% Tween 20) 3 times for 15 minutes each time. Ovaries were then blocked with PBTB (1X PBS, 0.2% Tween 20, 5% fetal bovine serum) for 1 hour at room temperature. The ovaries were treated with primary antibodies in PBTB at 4°C overnight. The following primary antibodies were used for immunostaining. Developmental Studies Hybridoma Bank (DSHB): mouse anti-Armadillo (N27A1, 1:200, concentrate), mouse anti-discs large (4F3, 1:200, concentrate), mouse anti-Notch extracellular domain (C458.2H, 1:200, concentrate), mouse anti-Cut (2B10, 1:400, concentrate), mouse anti-CycB (F2F4, 1:400, concentrate), mouse anti-Dynein heavy chain (2C11-2, 1:200, concentrate), mouse anti-Gurken (1D12, 1:200, concentrate). The primary antibodies were washed with PBST 3 times 15 min and then blocked with PBTB for 1 hr at room temperature. Ovaries were incubated with Alexafluor-conjugated secondary antibodies (1:500, Life technologies) in PBTB at 4°C overnight. Phalloidin 555 (1:50, Molecular Probes) was used to stain F-actin. Ovaries were washed with PBST for 15 minutes before staining nuclei with TO-PRO-3 (Life technologies, 1:1000) in PBST for 15 minutes. Ovaries were mounted in Vectashield (Vector laboratories). For Crumbs staining, Ovaries were dissected in PBS (Phosphate buffer saline) and fixed with boiled 8% paraformaldehyde in PBS and heptane (6:1) for 10 minutes. Samples were treated with heptane and methanol (1:2) for 30 seconds. They were then washed in methanol for 10 minutes. The ovaries were washed with PBST (1x PBS, 0.2% Tween 20) 2 times for 15 minutes each time. Ovaries were then blocked with PBTB (1x PBS, 0.2% Tween 20, 5% fetal bovine serum) for 30 minutes at room temperature. The ovaries were treated with mouse anti-Crumbs (Cq4, 1:100, concentrate, DSHB) in PBTB at 4°C overnight.

### Image acquisition and analysis of fixed samples

Images were taken on a confocal microscope (LSM710 or LSM780, Carl Zeiss) using 20x/0.5NA air objectives. Three laser lines were used based on the excitation of wavelength of the staining dyes which includes 488 nm, 561 nm and 633 nm wavelengths. Extent of migration (the migration index) was measured as a percentage of the distance travelled to the oocyte/nurse cell boundary in stage 10 egg chambers. ImageJ (https://imagej.nih.gov/ij/) was used for quantification of signal intensities in mosaic clusters using z-stack maximum projections. Raw integrated density was used as intensity values. For line scan profiles, maximum intensity images of Crumbs staining were generated in ImageJ. Background signal were subtracted. The plot profile function in ImageJ was used to measure signal intensities along lines drawn through the centre of border cell clusters and the peak analyser tool in OriginPro (Origin Lab) was used to calculate the area under peaks that were identified. The ratio of intensities at front, middle and back, were compared and normalised in Prism8 (Graphpad). Student’s t-tests statistical analysis were performed using Prism 8 (GraphPad): Figures were made using FigureApp in OMERO (Allan et al., 2012; Burel et al., 2015) and final assembly in Adobe Photoshop.

### Egg chamber culture and time-lapse imaging of live egg chambers

Live imaging of egg chamber culture was as previously described (Law et al., 2013; Prasad et al., 2007) with slight modification. Briefly, media for both dissection and live-imaging, comprised of Schneider media (Gibco), 15% fetal bovine serum, 0.1 mg/ml acidified insulin (Sigma), 9 μM FM4-64 dye (Molecular Probes) and 0.1 mg/ml Penstrep (Gibco) was freshly prepared. The pH of the media was adjusted to 6.90-6.95. Individual egg chambers from well fattened progeny of the right genotype were dissected and transferred to borosilicate glass bottom chambered coverglasses (ThermoFisher) for imaging. Imaging was done at 25^0^C. Time-lapse movies were acquired on an inverted confocal microscope (LSM 710; Carl Zeiss) using 20x/0.5NA air objectives. Two laser lines were used based on the excitation of wavelength of the endogenous GFP and FM4-64 dye, which are 488 nm, and 561 nm wavelengths respectively. 16-20 slices of Z-stacks were taken with 2.5 μm slices every 3 min.

## Genotypes of strains

**Figure 1**

(**A,E**) Control: *hsFLP; tub-Gal80, FRT40A/FRT40A; Act>CD2>Gal4,UAS-GFP/+*

(**B,F**) *Ataxin-7*^*HO3*^: *hsFLP; tub-Gal80, FRT40A/ Ataxin-7*^*HO3*^ *FRT40A; Act>CD2>Gal4,UAS-GFP/+*

(**C,D**) Quantification of (A,B).

(**G**) Quantification of (E,F).

**Figure 2**

(**A,G**) Control: *hsFLP; tub-Gal80, FRT40A/FRT40A; Act>CD2>Gal4,UAS-GFP/+*

(**B,C,H**) *Ataxin-7*^*HO3*^: *hsFLP; tub-Gal80, FRT40A/ Ataxin-7*^*HO3*^ *FRT40A; Act>CD2>Gal4,UAS-GFP/+*

(**D**) Quantification of (A,B).

(**E-F**) Quantification of (G-H).

**Figure 3**

(**A,D,F,H**) Control: *hsFLP; tub-Gal80, FRT40A/FRT40A; Act>CD2>Gal4,UAS-GFP/+*

(**B,E,G,I**) *Ataxin-7*^*HO3*^: *hsFLP; tub-Gal80, FRT40A/ Ataxin -7*^*HO3*^ *FRT40A; Act>CD2>Gal4,UAS-GFP/+*

(**C**) Quantification of (A,B and D,E)

**Figure 4**

(**A,C,E,G,I**) Control: *hsFLP; tub-Gal80, FRT40A/FRT40A; Act>CD2>Gal4,UAS-GFP/+*

(**B,D,F,H,J**) *Ataxin-7*^*HO3*^: *hsFLP; tub-Gal80, FRT40A/ Ataxin-7*^*HO3*^ *FRT40A; Act>CD2>Gal4,UAS-GFP/+*

**Figure 5**

(**A,B**) *not*^*IR*^: *hsFLP; Act>y>Gal4,UAS-GFP/ UAS-not*^*IR*^

(**C,D**) *ada2b*^*1*^: *hsFLP; Act>y>Gal4,UAS-GFP/+; FRT82B tubGAL80/ FRT82B Ada2B*

(**E,F**) *not*^*IR*^ *ada2b*^*1*^: *hsFLP; Act>y>Gal4,UAS-GFP/ UAS-not*^*IR*^; *FRT82B tubGAL80/FRT82B Ada2B*

**(G)** *Ataxin-7*^*HO3*^,*Notch: hsFLP; tub-Gal80, FRT40A/ Ataxin-7*^*HO3*^ *FRT40A; Act>CD2>Gal4,UAS-GFP/UAS-Notch*^*FL*^

(**H**) *Ataxin-7*^*HO3*^,*yki*^*IR*^: *hsFLP; tub-Gal80, FRT40A/ Ataxin-7*^*HO3*^ *FRT40A; Act>CD2>Gal4,UAS-GFP/UAS-yki*^*IR*^

(**I**) Quantification of (G,H)

## Movies

**Video 1: Time-lapse movie of normal follicular epithelium (10 frames/s) showing** normal monolayered architecture of the epithelium. MARCM clones are labelled with nuclear GFP; membrane is labelled with FM4-64 dye. Egg chamber genotype: *hsFLP; tub-Gal80, FRT40A/ FRT40A; Act>CD2>Gal4,UAS-GFP/+*.

**Video 2: Time-lapse movie of follicular epithelium (10 frames/s) showing** abnormal multi-layered epithelial architecture. MARCM clones are labelled with nuclear GFP; membrane is labelled with FM4-64 dye. Egg chamber genotype: *hsFLP; tub-Gal80, FRT40A/ Ataxin-7*^*HO3*^ *FRT40A; Act>CD2>Gal4,UAS-GFP/+*.

## Acknowledgments

We thank the Developmental Studies Hybridoma Bank (DSHB) for antibodies, the Bloomington Stock Center, Susan Abmayr, Jerry Workman and Nic Tapon for fly stocks. Thanks also to the Liverpool Centre for Cell Imaging for support with microscopy and image analysis. The work was funded by the MRC (MR/K015931/1), NWCR (CR847) and a University of Liverpool international PhD fees waiver.

